# Functional marker CAPS-799 of the *TaPHT1;9-4B* gene is useful for screening phosphorus-efficient wheat cultivars

**DOI:** 10.1101/2024.02.23.581827

**Authors:** Jinfeng Wang, Zedong Chen, Huanting Shi, Chuang Lou, Kaixia Fu, Yaxin Wang, Bo Yu, Tiancai Guo, Yonghua Wang, Pengfei Wang, Guozhang Kang

## Abstract

**Context:** In our previous study, *TaPHT1;9-4B*, one key high-affinity Pi transporter, was found to greatly contribute to Pi acquisition and transportation, and its functional marker CAPS-799 was subsequently developed to identify its Pi-efficient elite haplotype.

**Objective:** The study aimed to screen a varieties of wheat cultivars by using the above CAPS-799, identify its Pi-efficient elite haplotype cultivars, and reveal its physiological mechanism.

**Methods:** Successive two-year field experiments without Pi fertilizer supply, and hydroponic experiment with low Pi (10 μM) were performed. P concentrations, biomasses, grain yields, yield components, root growth parameters, and *TaPHT1;9-4B* transcript levels were measured. Total P accumulation and transport efficiency, and the relative growth rates were calculated.

**Results:** Eight Pi-efficient wheat cultivars (*Hap3* haplotype) were screened out by using the CAPS-799 from 80 modern major cultivars, and in successive two-year field experiments, their grain yields, spike numbers, P absorption and transport efficiencies were significantly higher than those of *No*n-*Hap3* haplotypes (*Hap1, 2* and *4*) under no Pi fertilizer supply condition, and therefore, these eight cultivars belonged to Pi-efficient elite haplotype. *TaPHT1;9-4B* transcript levels in roots at the early stage of grain filling period in field experiment, and the relative growth rates of total root surface areas, volumes and mean root diameters of *Hap3* cultivars in hydroponic experiment, were markedly higher than other haplotypes.

**Conclusions:** CAPS-799 was a useful functional marker for screening Pi-efficient wheat cultivars, and its Pi-efficient wheat cultivars were characterized with higher *TaPHT1;9-4B* transcript levels and more roots.

**Implications:** CAPS-799 will be used to screen or develop Pi-efficient wheat cultivars.

## 1. Introduction

Phosphorus (P) is an indispensable nutrient for plant metabolic and developmental processes. Appropriate application of phosphorus fertilizer is conducive to yield increase (Li et al., 2010; Han et al., 2014; Jagdeep et al., 2022; Gao et al., 2023). In pursuit of economic benefits, however, the overuse of phosphorus fertilizer is customary. It not only wastes resources but also leads to the accumulation and migration of excess phosphorus into groundwater, resulting in freshwater eutrophication (Ramaekers et al., 2010; Kalkhajeh et al., 2017; Schmieder et al., 2018). Since phosphorus in soil is prone to forming insoluble phosphates with other metallic elements, the utilization of phosphorus to plants in soils is limited (Zhang et al., 2001; Zhang et al., 2014). Overuse of phosphorus fertilizer often fails to be fully utilized, resulting in excessive phosphorus, which does not contribute to the improvement of phosphorus utilization efficiency (Li et al., 2010). Phosphorus is a non-renewable resource, and it is estimated that phosphorus resources will be insufficient by 2050 and exhausted by the end of the century (Vance et al., 2003).The utilization rate of phosphorus fertilizer for wheat in the current season is less than 20% (Rawal et al., 2022). It is crucial to screen out or develop wheat cultivars with high phosphorus efficiency for achieving sustainable agricultural development.

In the dynamic changes of the wheat genome, a large number of variations are rapidly generated, resulting in differences in quantitative gene dosage, forming the wheat cultivars with significant financial and nutritional importance (Dubcovsky et al., 2007). The variation of phosphorus efficiency among different wheat cultivars has already been reported (Ozturk et al., 2005; Korkmaz et al., 2009; Wei et al., 2020). Enhancing P efficiency can be achieved by improving P absorption efficiency, P transport efficiency or/and P utilization efficiency (Wang et al., 2010a; Wang et al., 2010b; Heuer et al., 2017; Zheng et al., 2023). It is time-consuming for breeding high-Pi-efficient cultivars by using traditional breeding methods (Wang et al., 2010c), and micro-methods such as genetic engineering or chromosome engineering of chromosomal displacement or ectopic placement are beneficial to achieve rapid breeding (Raghothama, 1999; Colmer et al., 2006). Moreover, the fully annotated reference genome in bread wheat accelerates the study of wheat genomics, especially in specific functional genes (IWGSC 2018).

Molecular marker-assisted selection has the characteristics of easy operation, stability, and high reliability, and, it can be utilized to analyze the genetic composition of individuals at the molecular level to select better genotypes (Torres et al., 2010). Cleaved amplified polymorphic sequences (CAPS) markers are commonly used to detect SNP sites in wheat breeding because of their specificity, simple operation and low cost (Collard et al., 2005; Dean et al., 2012; Shavrukov, 2016). At present, CAPS functional markers are widely used in wheat, maize, rice, soybean, and other crops (Zhang et al., 2010; Dissanayaka et al., 2014; Zhang et al., 2017; Ochar et al., 2022). In wheat, the development of functional markers provides theoretical basis for rapid and high-quality breeding, such as the genes *Rht-B1b* and *Rht-D1b*, *PPO*, and *TaCwi-A1* (Ellis et al., 2002; He et al., 2007; Ma et al., 2012). In addition, the molecular marker of *TaSRL1-4A* is valuable for the improvement of the root system, plant architecture, and yield (Zhuang et al., 2021). CAPS marker *TaGW-4B* is significantly correlated with 1000-grain weight (Duan et al., 2020). CAPS marker (*TaGS1a*-CAPS) showed a high association with mineral nutrient during seedling stage and grain size traits (Guo et al., 2013).

To our knowledge, however, wheat Pi-efficient CAPS markers have not been identified, mainly because of typically quantitative traits, tedious methods, expensive costs, vulnerable root systems, susceptible environmental influences, and complex and huge wheat genome (Cobb et al., 2013; Yuan et al., 2017; Yang et al., 2021; Xu et al., 2022). In our previous study, one high-affinity phosphate transporter protein, TaPHT1;9-4B, was identified using both isobaric tagging for relative and absolute quantification (iTRAQ) and parallel reaction monitoring (PRM) methods, and its important roles in Pi uptake and translocation in wheat were confirmed using barley stripe mosaic virus-virus induced gene-silencing (BSMV-VIGS), CRISPR/Cas9 editing and transgene methods. Nucleotide diversity of the gene encoding TaPHT1;9-4B was analyzed among 21 landraces and 41 modern cultivars, and nine SNPs in its promoter regions were found, forming four haplotypes (*Hap1*-*4*). Under hydroponic condition with low Pi (50 μM), *Hap3* exhibited higher phosphorus concentrations, and promoter activities and transcript levels of *TaPHT1;9-4B* than other haplotypes, implying that *Hap3* could be a Pi-efficient haplotype. Base on their SNPs, a functional marker CAPS-799 was developed to distinguish *Hap3* wheat cultivars (Wang et al., 2021). In this study, successive two-year field experiments without Pi fertilizer supply are designed, more modern major wheat cultivars are selected, and grain yields and phosphorus efficiency are calculated, to further identify whether *Hap3* belongs to Pi-efficient wheat cultivars. And its physiological mechanisms are further explored by measuring its transcript levels and root architecture under both field and hydroponic conditions.

## 2. Materials and methods

### 2.1. Functional marker screening

In our previous study, development of CAPS-799 marker was based on the two SNPs at positions 799 (C/G) and 796 (C/G) for different *TaPHT1;9-4B* promoter haplotypes, and *TaPHT1;9-4B* promoters of *Hap3* cannot be digested by the restriction endonuclease *Fnu*4HI, resulting in a single fragment (703 bp). However, there are two fragments (457 bp and 246 bp) in other haplotypes after their promoter are digested by *Fnu*4HI. Therefore, *Hap3* wheat cultivars of *TaPHT1;9-4B* are easily distinguished (Wang et al., 2021).

In this study, plant materials of the 80 modern major wheat cultivars developed in recent 20 years were selected and shown in Table 1. Their DNAs were extracted and *TaPHT1;9-4B* promoters were amplified using the above CAPS-799 primers. Primers are shown in Table S1. PCR amplicons were purified and recovered, and then digested using *Fnu*4HI. *TaPHT1;9-4B* haplotypes among wheat cultivars were divided based on the above method.

**Table 1.**
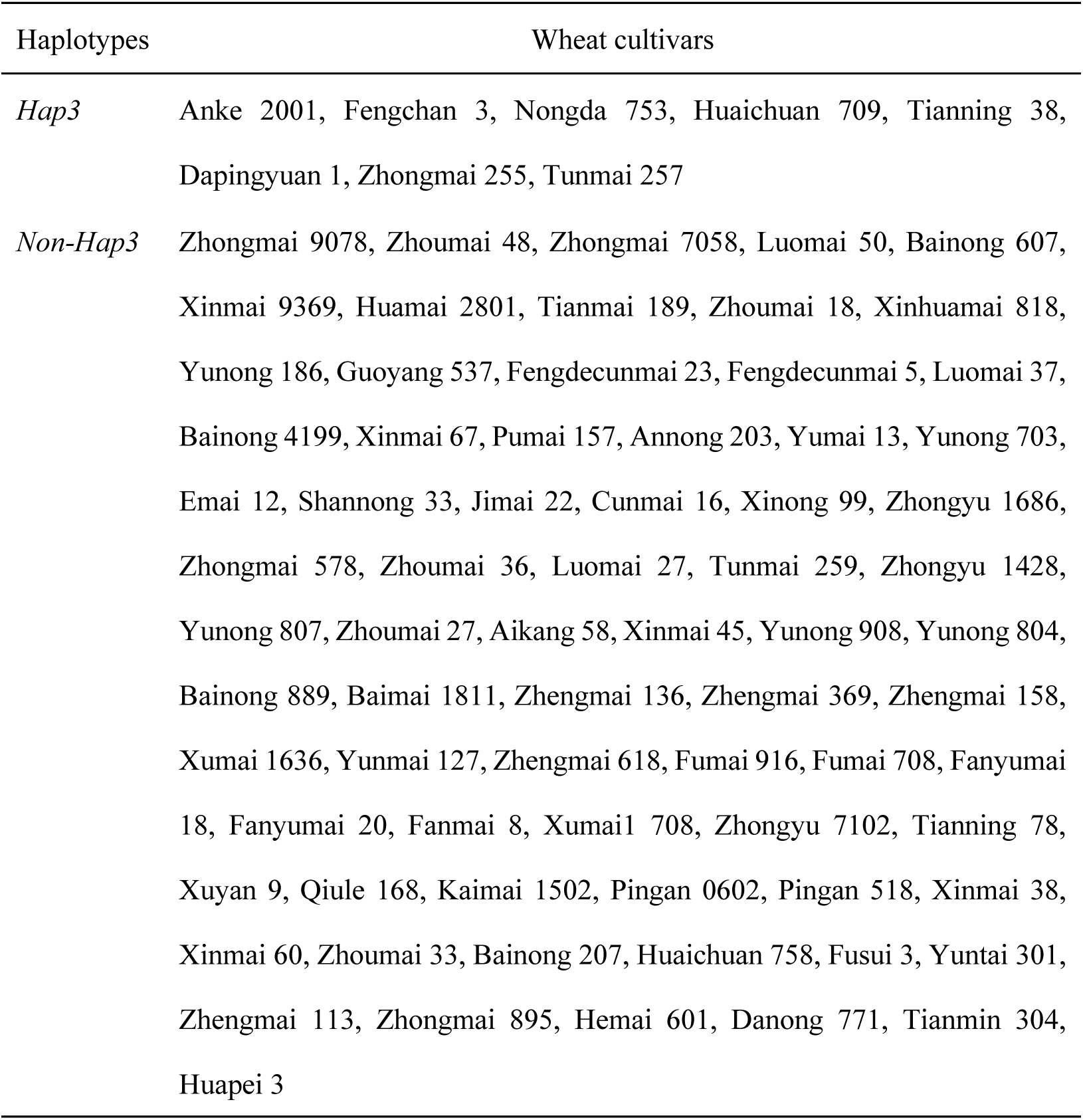
Haplotype types of wheat cultivars used in this study.

### 2.2. Field experiments

#### 2.2.1. Experimental design

Two-year successive field experiments were conducted at the Science and Education Park (35° 11 N, 113° 95 E) of Henan Agricultural University, Yuanyang County, Henan Province, China, and during the wheat growing seasons of 2021∼2022 and 2022∼2023. Soil at the top 0-20 cm and 20-40 cm layers were sampled before sowing, and the soil properties are shown in Table 2. Two Pi treatments were designed without Pi fertilizer (P0) supply and with normal Pi fertilizers (P120, 120 kg/ha P_2_O_5_) supply. Pure nitrogen 240 kg/ha and K_2_O 90 kg/ha were applied in the whole growth period of wheat in all plots. Phosphorus and potassium fertilizers were applied at one time before sowing. Half of nitrogen fertilizers were applied before sowing and the remained amounts were applied at jointing stage. Each wheat cultivar was separately planted in normal Pi-supply and no Pi plots (4.5 m × 0.8 m), and seeds of each wheat cultivar were sowed in 4 rows with 3 cm space between two adjacent seeds. Other field managements were carried out according to high-yield wheat practices.

**Table 2.** Soil properties in field experiments.

| Treatments | Layers | pH | Organic<br>matter<br>(%) | Total<br>nitrogen<br>(%) | Total<br>phosphorus<br>(%) | Total<br>potassium<br>(%) | Alkali-<br>hydrolyzed<br>nitrogen (ppm) | Available<br>phosphorus<br>(ppm) | Available<br>potassium<br>(ppm) |
| --- | --- | --- | --- | --- | --- | --- | --- | --- | --- |
| P120 | 0-20 | 8.24 | 1.08 | 0.12 | 0.07 | 0.88 | 90.77 | 24.34 | 110.93 |
|  | 20-40 | 8.38 | 0.51 | 0.11 | 0.06 | 0.99 | 69.22 | 21.70 | 101.15 |
| P0 | 0-20 | 8.39 | 1.21 | 0.10 | 0.07 | 0.95 | 93.36 | 9.14 | 142.36 |
|  | 20-40 | 8.56 | 0.66 | 0.07 | 0.06 | 0.91 | 66.63 | 8.00 | 147.13 |

#### 2.2.2. Sampling and measurements

Three plants of each wheat cultivar from P0 and P120 treatments were separately harvested at anthesis and maturity stages. Their roots were removed and the remaining parts were divided into two parts: the aboveground vegetative organs (stems, leaves, spikes, and sheaths) and the reproductive organ (grains). Plant samples were dried at 105℃ for 30 minutes and 80℃ to constant weight, and then dry weights were measured. Dried samples were pulverized and boiled with H_2_SO_4_-H_2_O_2_. P concentrations were measured using molybdate-blue colorimetric method (Chen et al., 2007). P accumulation (mg), P transport efficiency (%) and P utilization efficiency (g/g) were calculated as follows (Deng et al., 2018; Rezakhani et al., 2019; Zheng et al., 2022). Their relative growth rates (%), which represent the low phosphorus tolerance, were further calculated (Zhang et al., 2008).

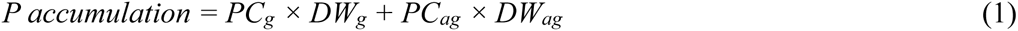

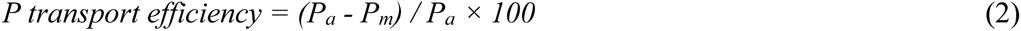

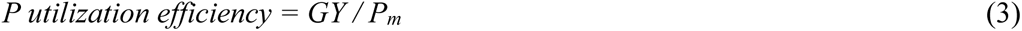

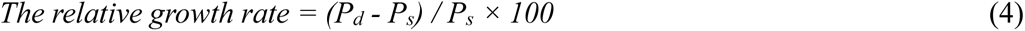

Where *PC_g_* and *PC_ag_* separately represent P concentration in grain and above-ground (%). *DW_g_* and *DW_ag_* represent dry weight in grain and above-ground (mg), respectively. *P_a_* and *P_m_* are the P accumulation at anthesis and maturity stages, respectively (mg). *GY* is the grain yields. *P_d_* and *P_s_* are the parameters in P0 and P120 treatments, respectively.

#### 2.2.3. Grain yields and components

At 10 days before maturity, spike numbers of each wheat cultivar were counted in three one-meter double rows, and their grain numbers per spike were calculated from randomly chosen twenty spikes. At maturity stage, grains in the above three one-meter double row plants were harvested and dried in the sun, and their 1000-grain weights and grain yields were separately weighted.

### 2.3. Hydroponic experiments

#### 2.3.1. Materials and design

Plant materials are listed in Table S2. Wheat seeds were disinfected with 75% ethanol and then rinsed for three times. Sterilized seeds were placed on glass dishes in a light incubator for hydroculture with 24℃/16℃ day/night temperature, 16 h/8 h day/night illumination duration, and 75% relative humidity. At the three-leaf stage after germination, the remained seeds were removed from seedlings to absorb nutrient from external solutions for supporting their autotrophic growth (Li et al., 2017). Then, seedlings were transferred to the Hoagland solution with normal Pi (200 μmol/L) and low Pi (10 μmol/L) for 21 d (Secco et al., 2010). Nutrient solutions were replaced every 2 days, and pH was kept among 5.5∼6.0.

#### 2.3.2. Sampling and measurements

Plants were sampled at 21 days after low Pi treatment. Roots of three plants from each treatment were rinsed, and their integrities were carefully protected. WinRHIZO software (Regent Instruments Canada) was used to measure root morphological parameters such as total root lengths, total root surface areas, total root volumes and mean root diameters (Ren et al., 2012). Then, plants were divided into two parts: aboveground shoots and underground roots. Samples were dried, weighed and then pulverized. P concentrations were measured and P accumulation amounts were calculated according to the above-mentioned methods.

### 2.4. Transcript levels of the TaPHT1;9-4B gene

Surface roots, stems below the spike and flag leaves at 10 days after anthesis from wheat plants in field experiment were sampled from P0 and P120 treatments. At this stage, wheat grains are quickly filled and a large amount of nutrients including Pi are needed. Their RNAs were extracted for synthesizing cDNA using Tissue RNA Isolation Kit (Beibei Biotechnology Co. Ltd., Cat# 082002). Transcript levels of *TaPHT1;9-4B* were measured by using the quantitative real-time PCR (qRT-PCR), and calculated according to transcript levels of the *TaActin* gene (Wang et al., 2021). Primers are listed in Table S1.

### 2.5. Statistical analysis

At least three biological repeats were designed and their statistical analyses were performed using SPSS 23.0 (IBM, Montauk, NY, USA). Differences significance test were analyzed using the Duncan method at *P* < 0.05 level. Graphs were plotted using Origin 2021 (Origin Lab Corporation, Northampton, MA, USA).

## 3. Results

### 3.1. Screening TaPHT1;9-4B haplotypes by using CAPS-799

*TaPHT1;9-4B* promoters of 80 wheat cultivars were separately amplified by using CAPS-799 primers, and digested by using *Fnu*4HI. Our results showed that PCR amplified fragments from 8 wheat cultivars were not be digested by *Fnu*4HI and there appeared one single fragment (703 bp), indicating that they belonged to *Hap3*. And the amplicons from the remaining 72 cultivars were cleaved into two fragments (457 and 246 bp), being divided into *Non-Hap3* (*Hap1*, *Hap2*, or *Hap4*) (Table 1, Fig. 1).

**Fig. 1.**
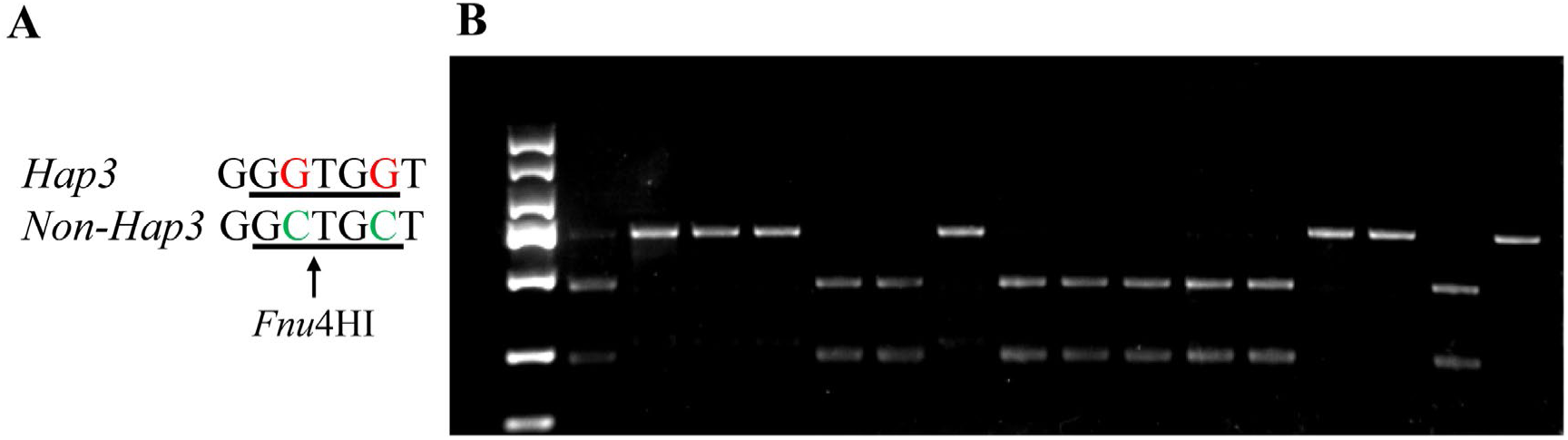
Electropherogram of representative *Hap3* wheat cultivars screened out by using CAPS-799. (A) The horizontal lines represent the base sequence recognized by the restriction endonuclease *Fnu*4HI. The arrow represents the specific cutting point of *Fnu*4HI. (B) *TaPHT1;9-4B* promoters were amplified by using CAPS-799 primers (Table S1) and digested by using *Fnu*4HI, and there appeared one single fragment (703 bp) in *Hap3* wheat cultivars, and two fragments (457 and 246 bp) in *Non-Hap3* (*Hap1/Hap2/Hap4*) wheat cultivars.

### 3.2. P uptake and utilization efficiencies of TaPHT1;9-4B haplotypes in field

The above *Hap3* and *Non*-*Hap3* wheat cultivars (Table 1), were grown for successive two-year field experiments to compare their phosphorus efficiencies. Total P accumulations and P utilization efficiencies reflect the total P, which is absorbed by roots and transported to the above-ground organs (P uptake efficiency), and the ratio of grain yields to total P, which is transported from other organs and utilized in grains, respectively. Our results showed that, under sufficient Pi supply condition (P120), there were insignificant differences in P accumulation between *Hap3* and *Non-Hap3* cultivars in successive two-year field experiments (Fig. 2A and C, left) and P utilization efficiency in 2022∼2023 wheat growth season (Fig. 2D, left), although significant difference appeared in P utilization efficiencies between *Hap3* and *Non-Hap3* cultivars in 2021∼2022 wheat growth season (Fig. 2B, left).

**Fig. 2.**
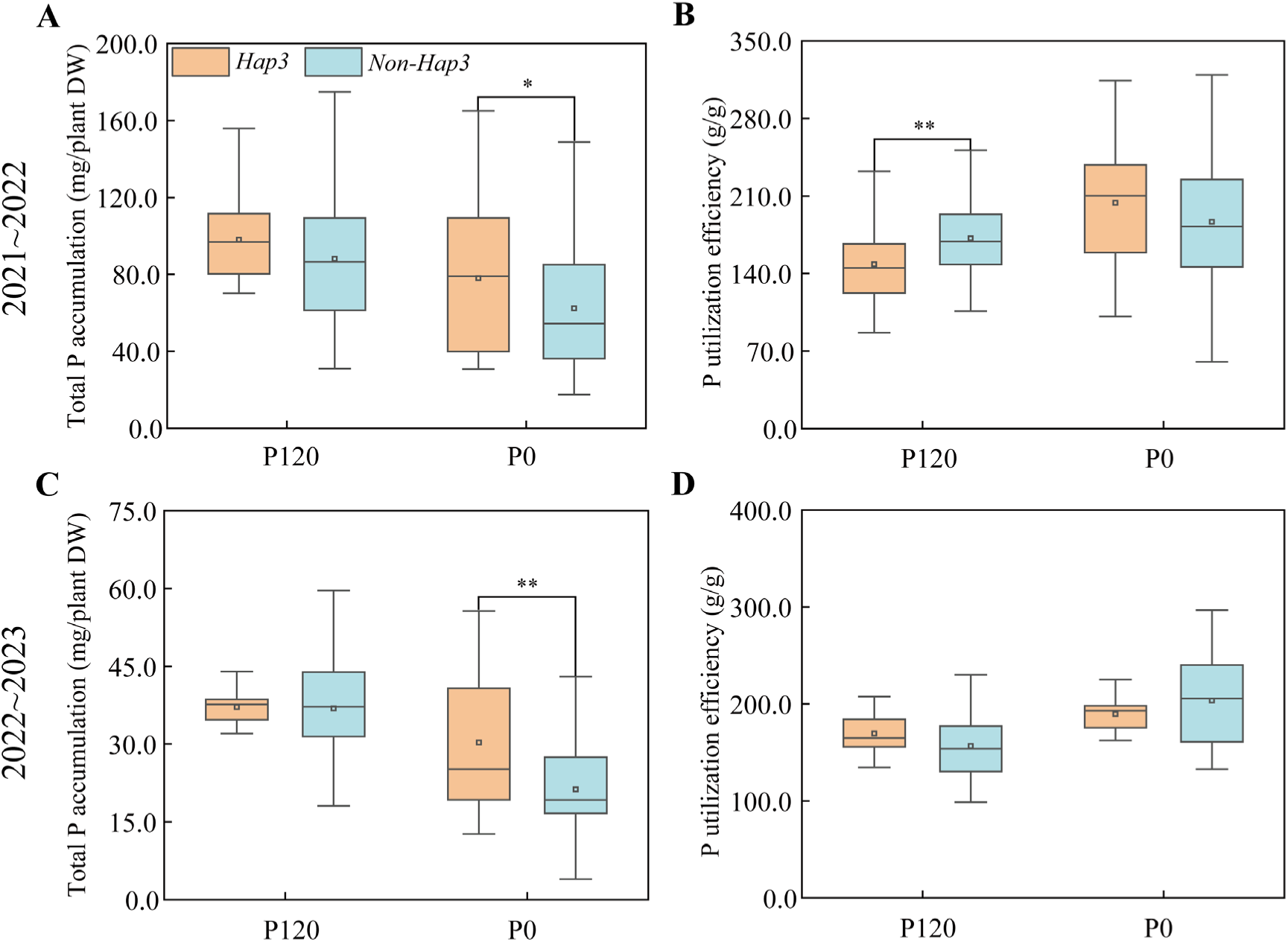
Phosphorous accumulation and utilization efficiencies of *Hap3* and *Non-Hap3* wheat cultivars. (A, B) Phosphorous accumulation and utilization efficiencies in field experiments in 2021∼2022. (C, D) Phosphorous accumulation and utilization efficiencies in field experiments in 2022∼2023. P120, normal phosphorus fertilizer application with 120 kg/ha P_2_O_5_; P0, no phosphorus fertilizer application. The asterisks indicate statistically significant differences (*, *P* < 0.05; **, *P* < 0.01; ***, *P* < 0.001; Duncan test).

Under no external Pi supply condition (P0), however, total P accumulation of *Hap3* wheat cultivars were significantly higher (+33.80%) than those of *Non-Hap3* cultivars (Fig. 2A and C, right). The relative growth rates of total P accumulation were further calculated. And similar results appeared in *Hap3* and *Non*-*Hap3* wheat cultivars (Fig. S1A, C). Under P0 condition, there were insignificant differences in P utilization efficiencies between *Hap3* and *Non*-*Hap3* wheat cultivars (Fig. 2B and D, right).

### 3.3. P transport efficiency of TaPHT1;9-4B haplotypes in field

The aboveground vegetative organs (stems, leaves, spikes, and sheaths) and the reproductive organ (grains) of both *Hap3* and *Non*-*Hap3* wheat cultivars at anthesis and maturity stages were separately harvested to measure its biomasses and P concentrations and further calculate their P transport efficiencies, which reflect the P accumulation transferred from vegetative organs to reproductive organ. Our results showed that, under P120 condition, P transport efficiencies of *Hap3* wheat cultivars were higher than *Non-Hap3*, insignificantly in 2021∼2022 and significantly in 2022∼2023 (Fig. 3A and B, left). Under P0 condition, however, this parameter in *Hap3* was remarkably higher (+13.84%) than *Non*-*Hap3* in successive two-year field experiments (Fig. 3A and B, right). And their relative growth rates of *Hap3* cultivars were also obviously higher in 2021∼2022 (Fig. S2A).

**Fig. 3.**
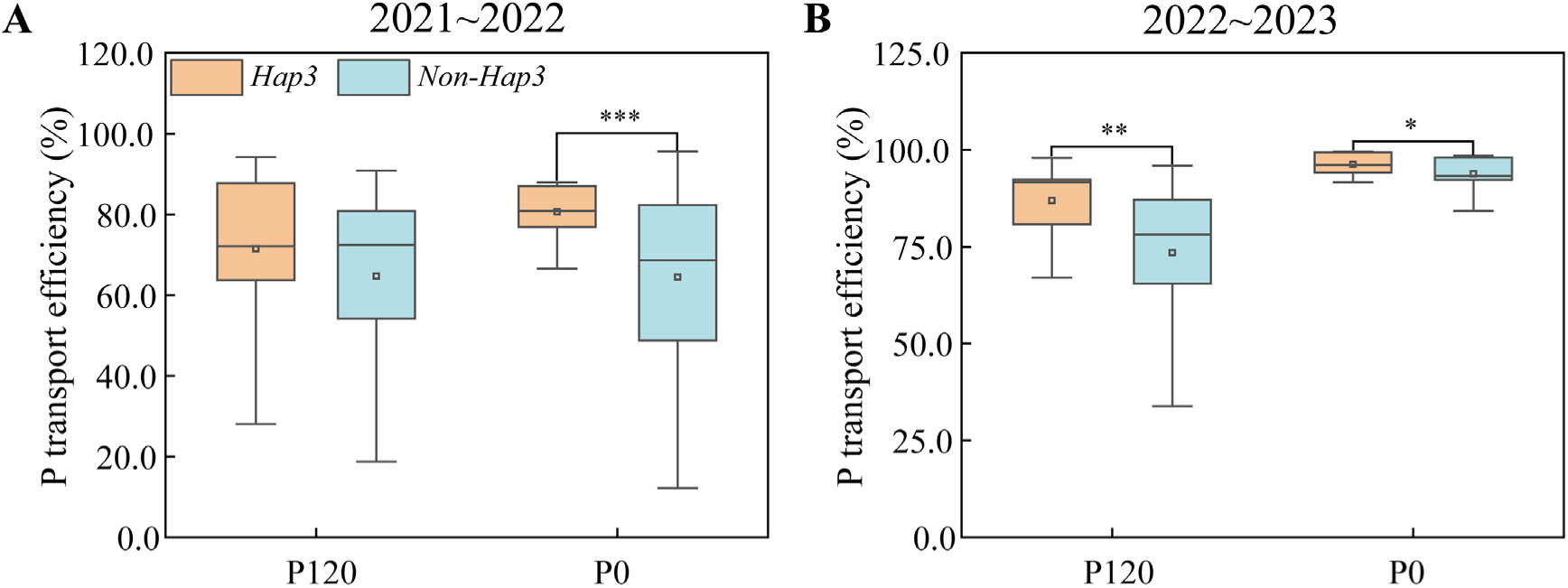
Phosphorous transport efficiencies of *Hap3* and *Non-Hap3* wheat cultivars. (A, B) Phosphorous transport efficiencies in field experiments in 2021∼2022 and 2022∼2023, respectively. P120, P0, and asterisks were annotated as same in Fig. 2.

### 3.4. Grain yields and components of TaPHT1;9-4B haplotypes in field

After harvesting, grain yields and their components were also compared between *Hap3* and *Non*-*Hap3* wheat cultivars. Under P120 condition, similarly, there were insignificant differences in grain yields between *Hap3* and *Non-Hap3* in successive two-year field experiments (Fig. 4A and E, left). Under P0 condition, however, grain yields in the former were remarkably higher than the latter, by +20.89% in 2021∼2022 and +26.92% in 2022∼2023, respectively (Fig. 4A and E, right). The relative growth rates of their grain yields were further compared, and our data indicated that, compared with *Non-Hap3*, this parameter in *Hap3* cultivars significantly increased by +54.71% in 2021-2023 (Fig. S3A, E).

**Fig. 4.**
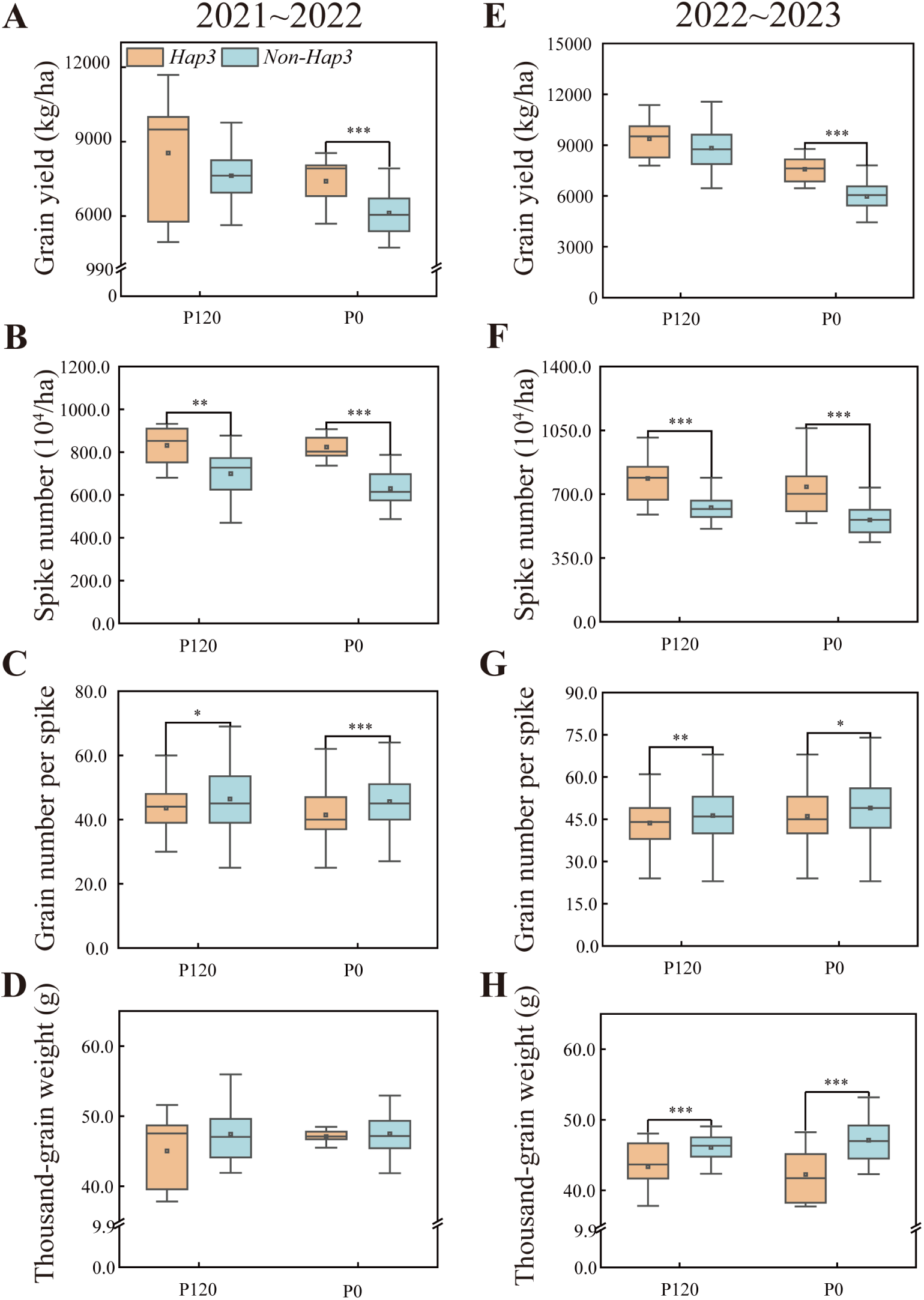
Grain yields of *Hap3* and *Non-Hap3* wheat cultivars. (A-D) Grain yields and components in field experiments in 2021∼2022. (E-H) Grain yields and components in field experiments in 2022∼2023. P120, P0, and asterisks were annotated as same in Fig. 2.

Further analysis of the yield components revealed that, under P0 condition, spike numbers in *Hap3* cultivars were significantly higher than those of *Non-Hap3* cultivars, by +30.79% in 2021∼2022 and +32.53% in 2022∼2023, respectively (Fig. 4B and F, right). The spike number relative growth rates were further calculated, and similarly, this parameter in *Hap3* was higher than *Non-Hap3*, by +44.05% in 2021∼2022 and +67.95% in 2022∼2023, respectively (Fig. S3B, F). Other two components (grain numbers per spike and thousand-grain weights) in *Hap3* cultivars were markedly or slightly lower than those in *Non-Hap3* cultivars in successive two-year field experiments (Fig. 4C, D, G, and H, right). These suggested that, under P0 condition, increased grain yields in *Hap3* cultivars were mainly from more spike numbers. Under P120 condition, three components of grain yields in both *Hap3* and *Non*-*Hap3* wheat cultivars exhibited the similar increased or decreased tendencies (Fig. 4B-D, and F-H, left), resulting in their similar grain yields (Fig. 4A and E, left).

### 3.5. Transcript levels of the TaPHT1;9-4B gene in field

Under field soil condition (P120 and P0), transcript levels of the *TaPHT1;9-4B* gene in roots of both *Hap3* and *Non*-*Hap3* wheat cultivars were remarkable higher by 50 and 6 times than those in stems and leaves, respectively (Fig. 5). Transcript levels of *TaPHT1;9-4B* were very low in stems and leaves of *Hap3* and *Non*-*Hap3* wheat cultivars either in P120 or P0 conditions (Fig. 5, middle and right). Under P120 condition, *TaPHT1;9-4B* transcript levels in roots of two haplotypes were insignificant between *Hap3* and *Non-Hap3*. Under P0 condition, however, transcript levels in roots of *Hap3* wheat cultivars were significantly greater (+100.74%) than *Non-Hap3* cultivars (Fig. 5, left).

**Fig. 5.**
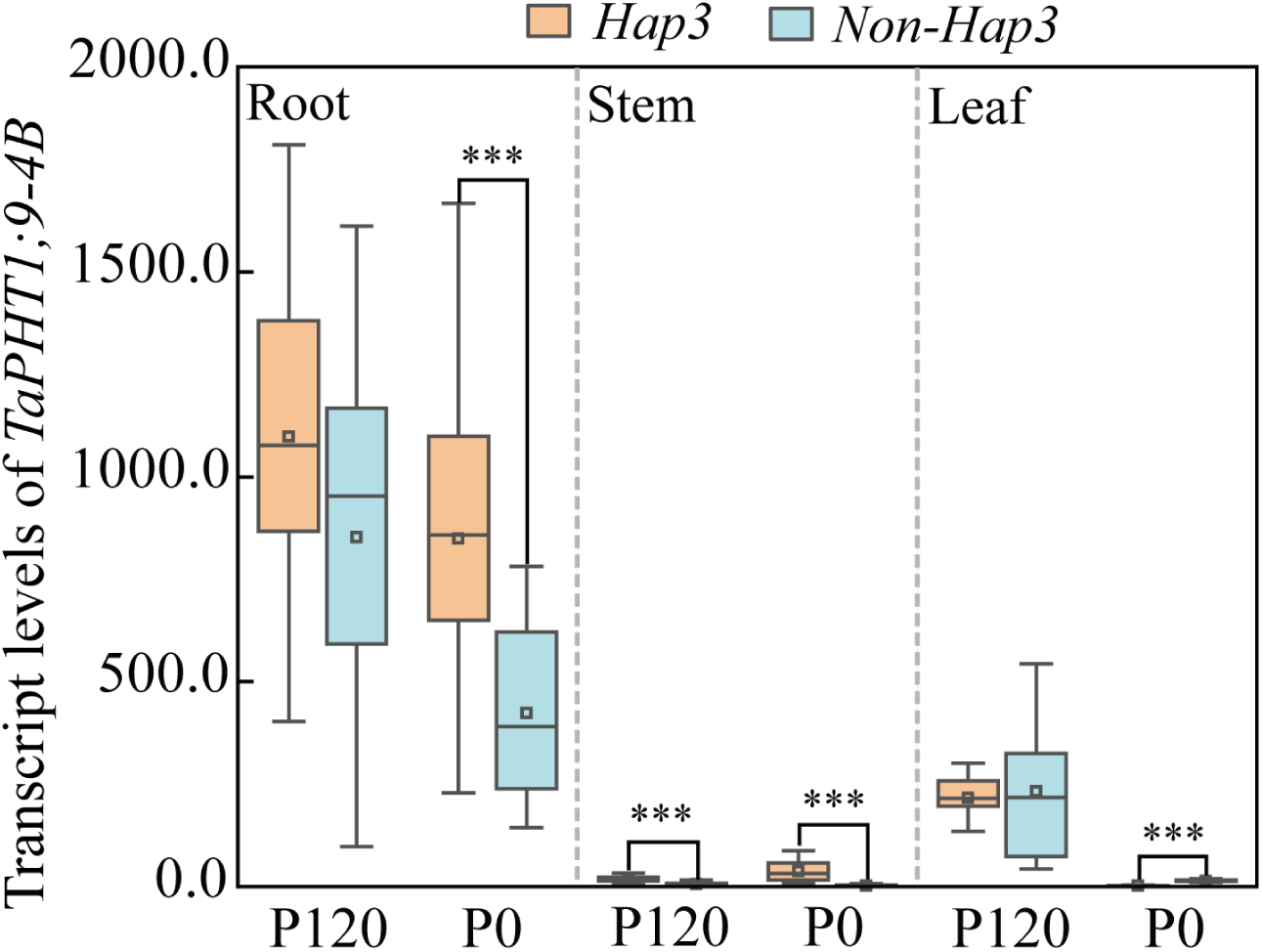
Transcript levels of *TaPHT1;9-4B* in root, stem and leaf tissues in field experiments at filling stage. *TaActin* is used as internal control gene. P120, P0, and asterisks were annotated as same in Fig. 2.

### 3.6. Root architecture among TaPHT1;9-4B haplotypes in hydroponics solutions

To avoid the soil disturbance as possible, and precisely measure the root architecture and their associated parameters, a hydroponics experiment was performed on *Hap3* and *Non*-*Hap3* wheat cultivars. Seedlings of six *Hap3* wheat cultivars and 12 *Non*-*Hap3* wheat cultivars were suffered from low Pi stress for 21 d and photographed. Under low Pi stress, more and longer roots length occurred in *Hap3* cultivars (Fig. 6A). These qualitative phenotypic changes were further verified by quantitative analysis. The relative growth rates of root parameters (total root surface area, total root volume and mean root diameter) in *Hap3* cultivars were significantly greater than *Non-Hap3* wheat cultivars (Fig. 6E, F, and G), and the relative growth rates of root dry weight and P accumulation of *Hap3* wheat cultivars were remarkably higher than those in *Non-Hap3* cultivars (Fig. 6B and C).

**Fig. 6.**
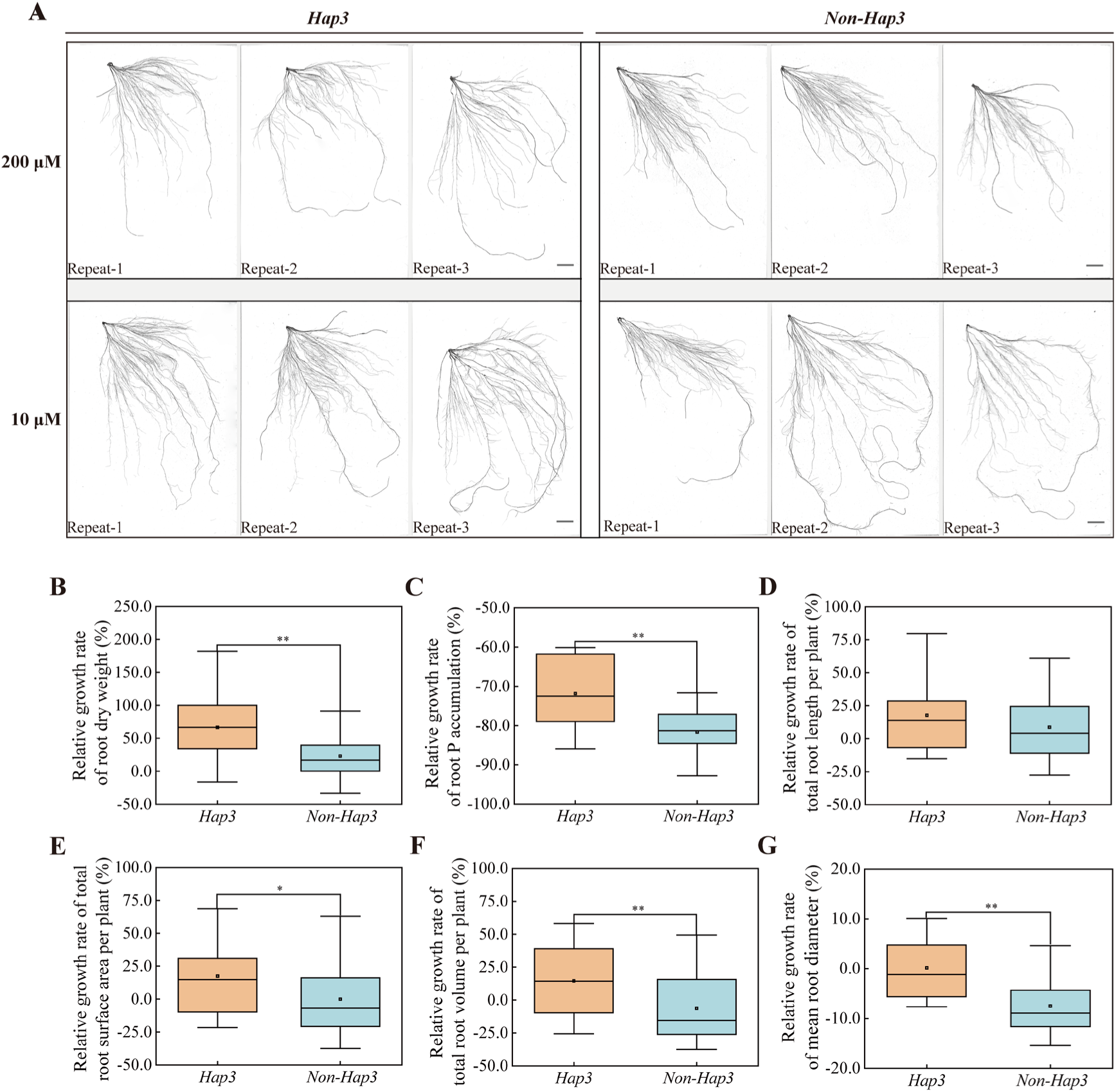
Root phenotypes and growth parameters of both *Hap3* and *Non-Hap3* wheat cultivars in hydroponics experiment. (A) Root scanning maps. Bar = 3 cm. 200 μM, normal phosphorus solution applied with 200 μmol/L Pi; 10 μM, low phosphorus solution applied with 10 μmol/L Pi. (B-G) The relative growth rates of root growth parameters. Asterisks were annotated as same in Fig. 2.

## 4. Discussion

### 4.1. CAPS-799 marker is useful for screening Pi-efficient wheat cultivars

Haplotype-based functional markers (etc., CAPS) are more reliably and easily used for applied breeding because they are derived from SNPs within the target genes or their regulatory sequences and are more likely to be related to phenotype variations (Roso et al., 2010; Ravikiran et al., 2018; Rasheed and Xia, 2019; Liu et al., 2022). In hydroponic solutions, in our previous study, *Hap3* of *TaPHT1;9-4B* were identified by CAPS-799 functional marker and speculated to be Pi-efficient wheat cultivars by evaluating the *TaPHT1;9-4B* expression levels, biomasses, and P contents (Wang et al., 2021). Compared to the hydroponic solutions, the soil Pi availabilities for growing plants are more complicated because of slow soil Pi mobility and low Pi availability (Grün et al., 2018). Under the field conditions, accordingly, this study further evaluated P efficiencies and yields of *TaPHT1;9-4B Hap3* and *Non*-*Hap3* by using 80 modern major cultivars and performing successive two-year field experiments with P120 (120 kg/ha P_2_O_5_ supply) and P0 (without Pi fertilizer supply) treatments. And more parameters (P efficiency, grain yield, etc.) and more growth stages (anthesis, grain filling and maturity) were measured. Our data indicated that, among the above selected wheat cultivars, eight were screened out by using CAPS-799 and they belonged to *Hap3* type (Table 1, Fig. 1). In subsequently successive two-year field experiments, total P accumulation, P transport efficiency, grain yield of eight *Hap3* cultivars under P0 treatment were significantly higher than *Non*-*Hap3* cultivars (Figs. 2-4). These further suggested that, compared to other three haplotypes, *Hap3* wheat cultivars were Pi-efficient, and CAPS-799 could be applicated practically as one key functional marker for screening *TaPHT1;9-4B Hap3*-mediated Pi-efficient wheat cultivars.

### 4.2. Possible Pi-efficient mechanisms on Hap3 wheat cultivars

PHTs are categorized into PHT1, PHT2, PHT3, and PHT4 subfamilies in plants. PHT2, PHT3, and PHT4 subfamilies comprise low-affinity Pi transporters, which mainly account for Pi acquisition when Pi supply is sufficient. Most of PHT1 members are low Pi-specific, whereas several PHT1s have important roles in response to both low and/or high Pi supply (Smith et al., 2003; Raghothama and Karthikeyan, 2005). Both OsPHT1;6/1;8 and HvPHT1;1 are responsive to Pi deficiency, whereas OsPHT1;2 and HvPHT1;6 are high Pi-responsive (Ai et al., 2009; Jia et al., 2011; Preuss et al.,2011; Rae et al., 2003). In the present study, obviously higher P uptake, utilization, and transport efficiencies, and grain yields in *TaPHT1;9-4B Hap3* appeared merely in P0 treatment, not in P120 (Figs. 2-4), implying that this transporter plays roles mainly in low Pi condition. Expression of plant *PHT1* members exhibited diverse patterns. For example, *OsPHT1;8* commonly expressed in stems, leaves, spikes and grains (Jia et al., 2017; Li et al., 2015), whereas the *OsPHT1;11/1;13* genes were root-specific (Paszkowski et al., 2002; Yang et al., 2012). Our results indicated that, the transcript levels of *TaPHT1;9-4B* richly expressed in roots (Fig. 5), indicating that it could mainly function in this organ.

Plant *PHT1s* roles are growth-stage specific (Römer and Schilling, 1986; Grün et al., 2018). *TaPHT1;1* in roots of field-derived wheat plants preferentially functioned at the beginning of stem elongation under Pi starved stress, whereas *TaPHT1;11* played roles mainly at both anthesis and milk ripening stages (Grün et al., 2018). In this study, the increased spike numbers of *TaPHT1;9-4B Hap3* wheat cultivars under P0 conditions gave rise to the obviously increased grain yields (Fig. 4), which were formed in field before the elongation stage. This implied that TaPHT1;9-4B mainly functioned in Pi uptake and transport at early growth stage.

Plant PHT1 members caused various root architecture changes (Maharajan et al., 2017; Zheng et al., 2021), and BnPHT1;4 under Pi-starved stress are responsive for longer primary roots and lower lateral root density (Ren et al., 2014). In this study, similarly, *Hap3* cultivars suffered from P0 stress also exhibited visible changes of root architecture, quantitatively confirmed by the relative higher growth rates of root parameters (Fig. 6).

A possible working model of the TaPHT1;9-4B protein in both the *Hap3* and *Non-Hap3* cultivars was proposed (Fig. 7).

**Fig. 7.**
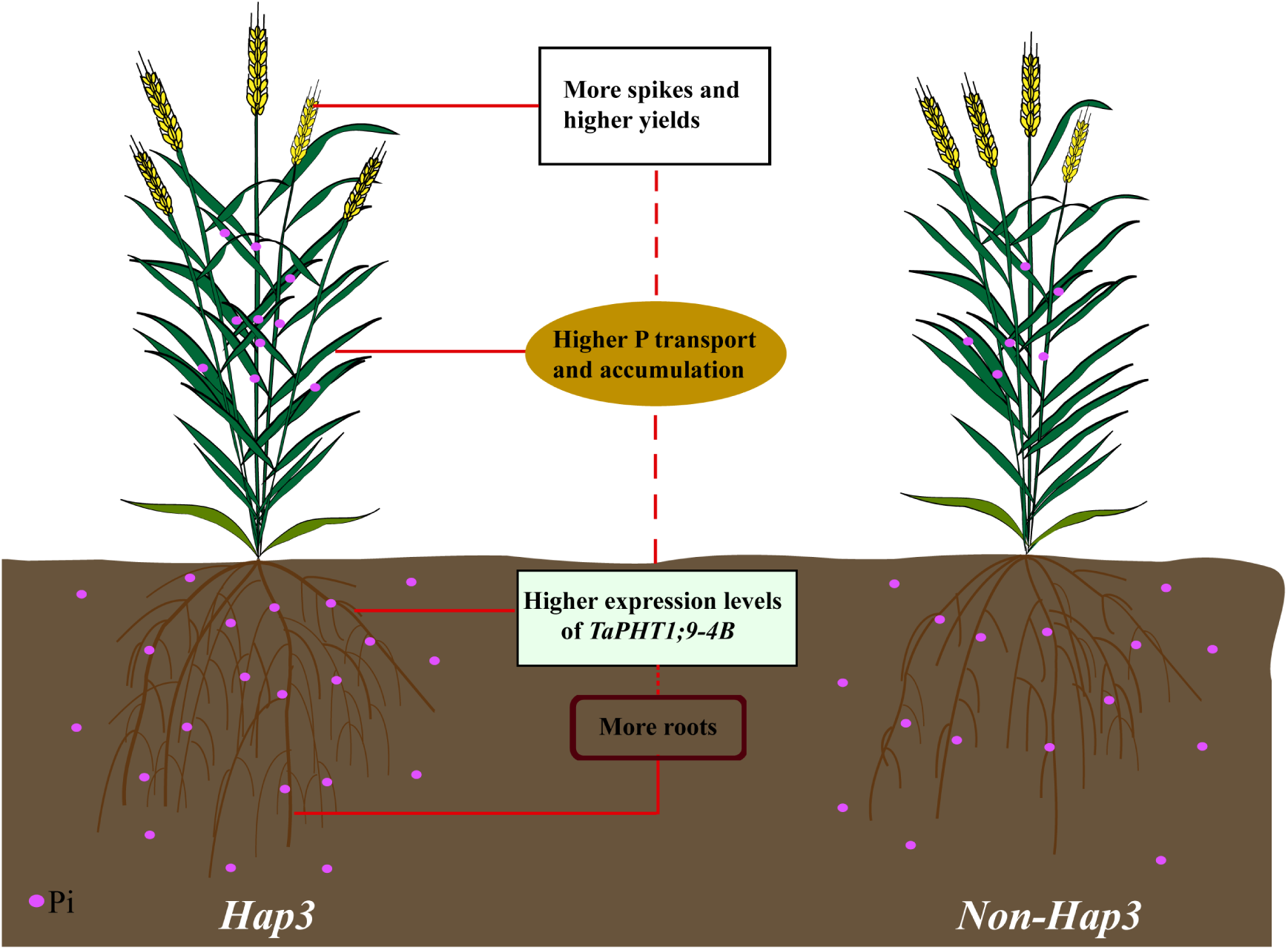
A working model of the TaPHT1;9-4B protein in both the *Hap3* and *Non-Hap3* wheat cultivars. Under low phosphorus condition, higher yields in *Hap3* cultivars could be due to more roots, higher expression levels of *TaPHT1;9-4B*, higher P uptake and transport efficiencies, and more spikes.

## 5. Conclusion

CAPS-799 was useful for screening Pi-efficient *Hap3* cultivars for the *TaPHT1;9-4B* gene, and this type of *Hap3* wheat cultivars under low Pi soil were characterized with higher P uptake and transport efficiencies, and grain yields. In *Hap3* wheat cultivars, *TaPHT1;9-4B* could mainly functioned in roots and at early growth stage.

## CRediT authorship contribution statement

**Guozhang Kang:** Conceptualization, Resources, Writing - Review & Editing, Project administration, Funding acquisition. **Pengfei Wang:** Methodology, Writing - Review & Editing, Supervision. **Jinfeng Wang:** Methodology, Validation, Formal analysis, Investigation, Writing - Original draft, Writing - Review & Editing, Visualization. **Zedong Chen:** Investigation, Writing - Review & Editing. **Huanting Shi, Chuang Lou, Kaixia Fu, and Yaxin Wang:** Investigation. **Bo Yu:** Writing - Review & Editing. **Tiancai Guo:** Supervision, Project administration. **Yonghua Wang:** Methodology, Resources, Supervision.

## Data Availability

Data will be made available on request.

## Acknowledgements

The work was financially supported by the Projects of Science and Technology of Henan Province (222102110039 and 232102111104), the China Agriculture Research System of MOF and MARA (CARS-03), and Shennong Laboratory of Henan Province (SN01-2022-01).

---

**Table S1.**
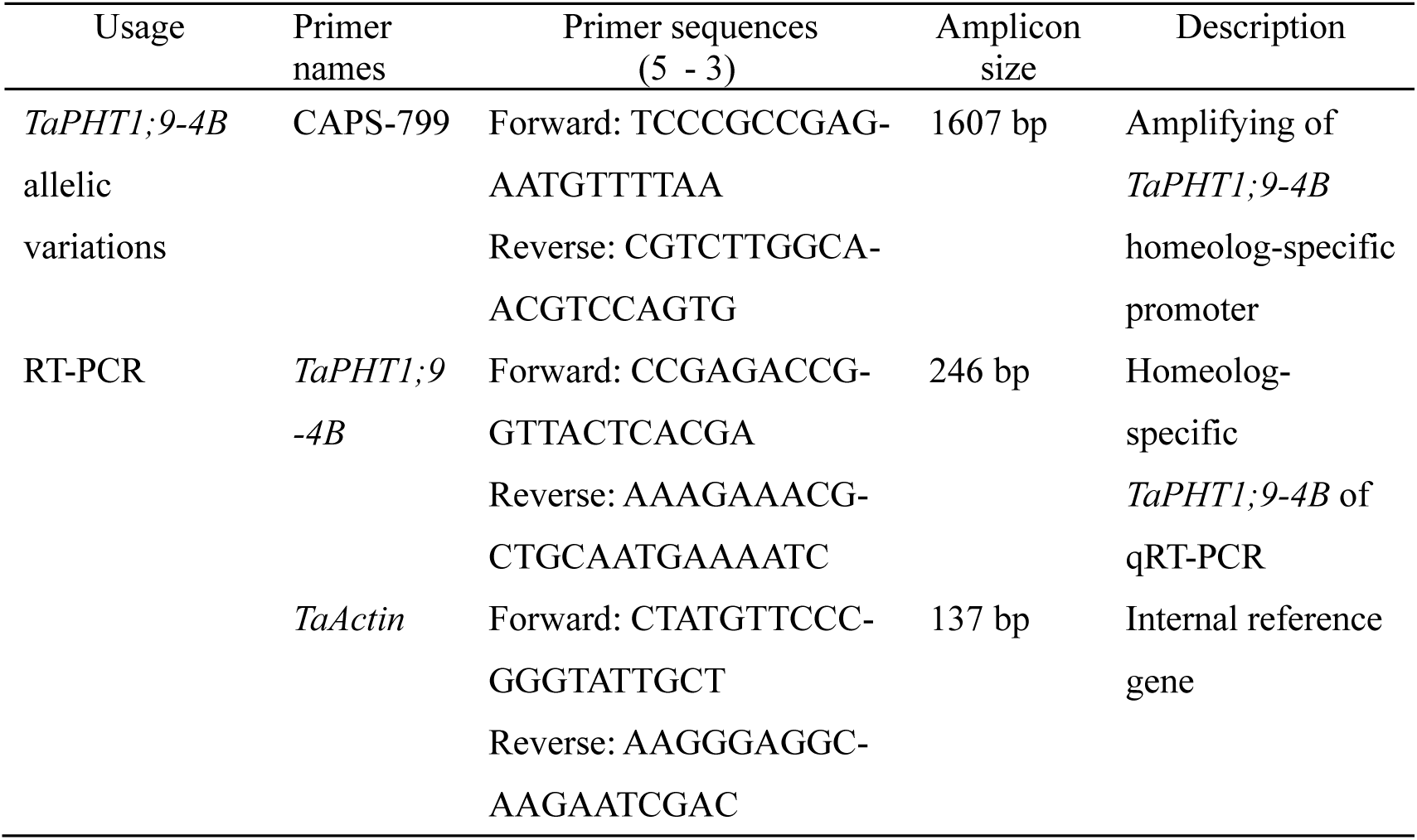
Primers used in this study.

## References

Ai, P.H., Sun, S.B., Zhao, J.N., Fan, X.R., Xin, W.J., Guo, Q., Yu, L., Shen, Q.R., Wu, P., Miller, A.J., Xu, G.H., 2009. Two rice phosphate transporters, OsPht1;2 and OsPht1;6, have different functions and kinetic properties in uptake and translocation. Plant J. 57 (5), 798–809.

Chen, A.Q., Hu, J., Sun, S.B., Xu, G.H., 2007. Conservation and divergence of both phosphate- and mycorrhiza-regulated physiological responses and expression patterns of phosphate transporters in solanaceous species. New Phytol. 173 (4), 817–831.

Cobb, J.N., Declerck, G., Greenberg, A., Clark, R., McCouch, S., 2013. Next-generation phenotyping: requirements and strategies for enhancing our understanding of genotype-phenotype relationships and its relevance to crop improvement. Theor Appl Genet. 126 (4), 867–887.

Collard, B.C.Y., Jahufer, M.Z.Z., Brouwer, J.B., Pang, E.C.K., 2005. An introduction to markers, quantitative trait loci (QTL) mapping and marker-assisted selection for crop improvement: The basic concepts. Euphytica 142, 169–196.

Colmer, T.D., Flowers, T.J., Munns, R., 2006. Use of wild relatives to improve salt tolerance in wheat. J Exp Bot. 57 (5), 1059–1078.

Dean, R., Kan, J.A.L.V., Pretorius, Z.A., Hammond-Kosack, K.E., Pietro, A.D., Spanu, P.D., Rudd, J.J., Dickman, M., Kahmann, R., Ellis, J., Foster, G.D., 2012. The Top 10 fungal pathogens in molecular plant pathology. Mol Plant Pathol. 13 (4), 414–430.

Deng, Y., Teng, W., Tong, Y.P., Chen, X.P., Zou, C.Q., 2018. Phosphorus efficiency mechanisms of two wheat cultivars as affected by a range of phosphorus levels in the field. Front Plant Sci. 6 (9), 1614.

Dissanayaka, S., Kottearachchi, N.S., Weerasena, J., Peiris, M., 2014. Development of a CAPS marker for the *badh2.7* allele in Sri Lankan fragrant rice (*Oryza sativa*). Plant Breed. 133 (5), 560–565.

Duan, X.X., Yu, H.X., Ma, W.J., Sun, J.Q., Zhao, Y., Yang, R.C., Ning, T.Y., Li, Q.F., Liu, Q.Q., Guo, T.T., Yan, M., Tian, J.C., Chen, J.S., 2020. A major and stable QTL controlling wheat thousand grain weight: identification, characterization, and CAPS marker development. Mol Breed. 40 (7), 68.

Dubcovsky, J., Dvorak, J., 2007. Genome plasticity a key factor in the success of polyploid wheat under domestication. Science 316 (5833), 1862–1866.

Ellis, H., Spielmeyer, W., Gale, R., Rebetzke, J., Richards, A., 2002. “Perfect” markers for the *Rht-B1b* and *Rht-D1b* dwarfing genes in wheat. Theor Appl Genet. 105 (6-7), 1038–1042.

Gao, Z.Y., Cao, H.B., Huang, M., Bao, M., Qiu, W.H., Liu, J.S., 2023. Winter wheat yield and soil critical phosphorus value response to yearly rainfall and P fertilization on the Loess Plateau of China. Field Crop Res. 296, 108921.

Grün, A., Buchner, P., Broadley, M.R., Hawkesford, M.J., 2018. Identification and expression profiling of Pht1 phosphate transporters in wheat in controlled environments and in the field. Plant Biol. 20 (2), 374–389.

Guo, Y., Sun, J.J., Zhang, G.Z., Wang, Y.Y., Kong, F.M., Zhao, Y., Li, S.S., 2013. Haplotype, molecular marker and phenotype effects associated with mineral nutrient and grain size traits of *TaGS1a* in wheat. Field Crop Res. 154, 119–125.

Han, Y.Y., Zhou, S., Chen, Y.H., Kong, X.Z., Xu, Y., Wang, W., 2014. The involvement of expansins in responses to phosphorus availability in wheat, and its potentials in improving phosphorus efficiency of plants. Plant Physiol Biochem. 78, 53–62.

He, X.Y., He, Z.H., Zhang, L.P., Sun, D.J., Morris, C.F., Fuerst, E.P., Xia, X.C., 2007. Allelic variation of polyphenol oxidase (PPO) genes located on chromosomes 2A and 2D and development of functional markers for the PPO genes in common wheat. Theor Appl Genet. 115 (1), 47–58.

Heuer, S., Gaxiola, R., Schilling, R., Herrera-Estrella, L., López-Arredondo, D., Wissuwa, M., Delhaize, E., Rouached, H., 2017. Improving phosphorus use efficiency: a complex trait with emerging opportunities. Plant J. 90 (5), 868–885.

Jagdeep, S., Brar, B.S., 2022. Build-up and utilization of phosphorus with continues fertilization in maize-wheat cropping sequence. Field Crop Res. 276, 108389.

Jia, H.F., Ren, H.Y., Gu, M., Zhao, J.N., Sun, S.B., Zhang, X., Chen, J.Y., Wu, P., Xu, G.H., 2011. The phosphate transporter gene *OsPht1;8* is involved in phosphate homeostasis in rice. Plant Physiol. 156 (3), 1164–1175.

Jia, H.F., Zhang, S.T., Wang, L.Z., Yang, Y.X., Zhang, H.Y., Cui, H., Shao, H.F., Xu, G.H., 2017. OsPht1;8, a phosphate transporter, is involved in auxin and phosphate starvation response in rice. J Exp Bot. 68 (18), 5057–5068.

Kalkhajeh, Y.K., Huang, B., Hu, W.Y., Holm, P.E., Hansen, H.C.B., 2017. Phosphorus saturation and mobilization in two typical Chinese greenhouse vegetable soils. Chemosphere 187, 440–441.

Korkmaz, K., Ibrikci, H., Karnez, E., Buyuk, G., Ryan, J., Ulger, A.C., Oguz, H., 2009. Phosphorus use efficiency of wheat genotypes grown in calcareous soils. J Plant Nutr. 32, 2094–2106.

Li, M., Guo, X.H., Zhang, M., Wang, X.P., Zhang, G.D., Tian, Y.C., Wang, Z.L., 2010. Mapping QTLs for grain yield and yield components under high and low phosphorus treatments in maize (*Zea mays* L.). Plant Sci. 178 (5), 454–462.

Li, Y.T., Zhang, J., Zhang, X., Fan, H.M., Gu, M., Qu, H.Y., Xu, G.H., 2015. Phosphate transporter OsPht1;8 in rice plays an important role in phosphorus redistribution from source to sink organs and allocation between embryo and endosperm of seeds. Plant Sci. 230, 23–32.

Li, G.Z., Wu, Y.F., Liu, G.Y., Xiao, X.H., Wang, P.F., Gao, T., Xu, M.J., Han, Q.X., Wang, Y.H., Guo, T.C., Kang, G.Z., 2017. Large-scale proteomics combined with transgenic experiments demonstrates an important role of jasmonic acid in potassium deficiency response in wheat and rice. Mol Cell Proteomics. 16 (11), 1889–1905.

Liu, X.J., Liu, X.C., Sun, H.G., Hao, C.Y., Wang, X.X., Rong, Z.J., Feng, Z.Y., 2022. Validation of CAPS marker WR003 for the leaf rust resistance gene *Lr1* and the molecular evolution of *Lr1* in wheat. Czech J Genet Plant Breed. 58 (4), 223–232.

Ma, D.Y., Yan, J., He, Z.H., Wu, L., Xia, X.C., 2012. Characterization of a cell wall invertase gene *TaCwi-A1* on common wheat chromosome 2A and development of functional markers. Mol Breed. 29 (1), 43–52.

Maharajan, T., Ceasar, S.A., Krishna, T.P.A., Ramakrishnan, M., Duraipandiyan, V., Abdulla, A.N., Ignacimuthu, S., 2017. Utilization of molecular markers for improving the phosphorus efficiency in crop plants. Plant Breed. 137 (1), 10–26.

Ochar, K., Su, B.H., Zhou, M.M., Liu, Z.X., Gao, H.W., F.Lamlom, S., Qiu, L.J., 2022. Identification of the genetic locus associated with the crinkled leaf phenotype in a soybean (*Glycine max* L.) mutant by BSA-Seq technology. J Integr Agric. 21 (12), 3524–3539.

Ozturk, L., Eker, S., Torun, B., Cakmak, I., 2005. Variation in phosphorus efficiency among 73 bread and durum wheat genotypes grown in a phosphorus-deficient calcareous soil. Plant Soil 269, 69–80.

Paszkowski, U., Kroken, S., Roux, C., Briggs, S.P., 2002. Rice phosphate transporters include an evolutionarily divergent gene specifically activated in arbuscular mycorrhizal symbiosis. Proc Natl Acad Sci U S A. 99 (20), 13324–13329.

Preuss, C.P., Huang, C.Y., Tyerman, S.D., 2011. Proton-coupled high-affinity phosphate transport revealed from heterologous characterization in *Xenopus* of barley-root plasma membrane transporter, HvPHT1;1. Plant Cell Environ. 34 (4), 681–689.

Rae, A.L., Cybinski, D.H., Jarmey, J.M., Smith, F.W., 2003. Characterization of two phosphate transporters from barley; evidence for diverse function and kinetic properties among members of the Pht1 family. Plant Mol Biol. 53, 27–36.

Raghothama, K.G., 1999. Phosphate acquisition. Annu Rev Plant Physiol Plant Mol Biol. 50, 665–693.

Raghothama, K.G., Karthikeyan, A.S., 2005. Phosphate acquisition. Plant Soil 274, 37–49.

Ramaekers, L., Remans, R., Rao, I.M., Blair, M.W., Vanderleyden, J., 2010. Strategies for improving phosphorus acquisition efficiency of crop plants. Field Crop Res. 117, 169–176.

Rasheed, A., Xia, X.C., 2019. From markers to genome-based breeding in wheat. Theor Appl Genet. 132 (3), 767–784.

Ravikiran, K.T., Krishnamurthy, S.L., Warraich, A.S., Sharma, P.C., 2018. Diversity and haplotypes of rice genotypes for seedling stage salinity tolerance analyzed through morpho-physiological and SSR markers. Field Crop Res. 220, 10–18.

Rawal, N., Pande, K.R., Shrestha, R., Vista, S.P., 2022. Nutrient use efficiency (NUE) of wheat (*Triticum aestivum* L.) as affected by NPK fertilization. PloS One 17, e0262771.

Ren, Y.Z., He, X., Liu, D.C., Li, J.J., Zhao, X.Q., Li, B., Tong, Y.P., Zhang, A.M., Li, Z.S., 2012. Major quantitative trait loci for seminal root morphology of wheat seedlings. Mol Breed. 30, 139–148.

Ren, F., Zhao, C.Z., Liu, C.S., Huang, K.L., Guo, Q.Q., Chang, L.L., Xiong, H., Li, X.B., 2014. A *Brassica napus* PHT1 phosphate transporter, BnPht1;4, promotes phosphate uptake and affects roots architecture of transgenic Arabidopsis. Plant Mol Biol. 86 (6), 595–607.

Rezakhani, L., Motesharezadeh, B., Tehrani, M.M., Etesami, H., Hosseini, H.M., 2019. Phosphate-solubilizing bacteria and silicon synergistically augment phosphorus (P) uptake by wheat (*Triticum aestivum* L.) plant fertilized with soluble or insoluble P source. Ecotoxicol Environ Saf. 173, 504–513.

Römer, R., Schilling, G., 1986. Phosphorus requirements of the wheat plant in various stages of its life cycle. Plant Soil 91, 221–229.

Roso, A.C., Merotto Jr, A., Delatorre, C.A., Menezes, V.G., 2010. Regional scale distribution of imidazolinone herbicide-resistant alleles in red rice (*Oryza sativa* L.) determined through SNP markers. Field Crop Res. 119, 175–182.

Schmieder, F., Bergstrom, L., Riddle, M., Gustafsson, J., Klysubun, W., Zehetner, F., Condron, L., Kirchmann, H., 2018. Phosphorus speciation in a long-term manure-amended soil profile - Evidence from wet chemical extraction, ^31^P-NMR and P K-edge XANES spectroscopy. Geoderma 322, 19–27.

Secco, D., Baumann, A., Poirier, Y., 2010. Characterization of the rice *PHO1* gene family reveals a key role for *OsPHO1;2* in phosphate homeostasis and the evolution of a distinct clade in dicotyledons. Plant Physiol. 152 (3), 1693–1704.

Shavrukov, Y., 2016. Comparison of SNP and CAPS markers application in genetic research in wheat and barley. BMC Plant Biol. 16, 11–15.

Smith, F.W., Mudge, S.R., Rae, A.L., Glassop, D., 2003. Phosphate transport in plants. Plant Soil 248, 71–83.

Torres, A.M., Avila, C.M., Gutierrez, N., Palomino, C., Moreno, M.T., Cubero, J.I., 2010. Marker-assisted selection in faba bean (*Vicia faba* L.). Field Crop Res. 115 (3), 243–252.

Vance, C.P., Uhde-Stone, C., Allan, D.L., 2003. Phosphorus acquisition and use: critical adaptations by plants for securing a nonrenewable resource. New Phytol. 157 (3), 423–447.

Wang, X.R., Shen, J.B., Liao, H., 2010a. Acquisition or utilization, which is more critical for enhancing phosphorus efficiency in modern crops? Plant Sci. 179 (4), 302–306.

Wang, L.Z., Chen, F.J., Zhang, F.S., Mi, G.H., 2010b. Two strategies for achieving higher yield under phosphorus deficiency in winter wheat grown in field conditions. Field Crop Res. 118, 36–42.

Wang, S.W., Yin, L.N., Tanaka, H., Tanaka, K., Tsujimoto, H., 2010c. Identification of wheat alien chromosome addition lines for breeding wheat with high phosphorus efficiency. Breed Sci. 60 (4), 371–379.

Wang, P.F., Li, G.Z., Li, G.W., Yuan, S.S., Wang, C.Y., Xie, Y.X., Guo, T.C., Kang, G.Z., Wang, D.W., 2021. TaPHT1;9-4B and its transcriptional regulator TaMYB4-7D contribute to phosphate uptake and plant growth in bread wheat. New Phytol. 231 (5), 1968–1983.

Wei, R.P., Wang, X., Zhang, W., Shen, J.N., Zhang, H.F., Gao, Y., Yang, L.Y., 2020. The improved phosphorus utilization and reduced phosphorus consumption of *ppk*-expressing transgenic rice. Field Crop Res. 248, 107715.

Xu, J.L., Gao, Z.Y., Liu, S., Elwafa, S.F.A., Tian, H., 2022. A multienvironmental evaluation of the N, P and K use efficiency of a large wheat diversity panel. Field Crop Res. 286, 108634.

Yang, S.Y., GrØnlund, M., Jakobsen, I., Grotemeyer, M.S., Rentsch, D., Miyao, A., Hirochika, H., Kumar, C.S., Sundaresan, V., Salamin, N., Catausan, S., Mattes, N., Heuer, S., Paszkowski, U., 2012. Nonredundant regulation of rice arbuscular mycorrhizal symbiosis by two members of the *PHOSPHATE TRANSPORTER1* gene family. Plant Cell 24 (10), 4236–4251.

Yang, M.J., Wang, C.R., Hassan, M.A., Li, F.J., Xia, X.C., Shi, S.B., Xiao, Y.G., He, Z.H., 2021. QTL mapping of root traits in wheat under different phosphorus levels using hydroponic culture. BMC Genomics. 22, 174.

Yuan, Y.Y., Gao, M.G., Zhang, M.X., Zheng, H.H., Zhou, X.W., Guo, Y., Zhao, Y., Kong, F.M., Li, S.S., 2017. QTL mapping for phosphorus efficiency and morphological traits at seedling and maturity stages in wheat. Front Plant Sci. 8, 614.

Zhang, M., Alva, A.K., Li, Y.C., Calvert, D.V., 2001. Aluminum and iron fractions affecting phosphorus solubility and reactions in selected sandy soils. Soil Sci. 166 (12), 940–948.

Zhang, H.W., Yu, H., Ye, X.S., Xu, F.S., 2008. Evaluation of phosphorus efficiency in rapeseed (*Brassica napus* L.) recombinant inbred lines at seedling stage. Acta Agron Sin. 34 (12), 2152–2159.

Zhang, Y., Xu, L., Zhang, D.F., Dai, J.R., Wang, S.C., 2010. Mapping of southern corn rust-resistant genes in the W2D inbred line of maize (*Zea mays* L.). Mol Breed. 25, 433–439.

Zhang, M., Li, C.L., Li, Y.C., Harris, W.G., 2014. Phosphate minerals and solubility in native and agricultural calcareous soils. Geoderma 232-234, 164–171.

Zhang, P.F., He, Z.H., Tian, X.L., Gao, F.M., Xu, D.A., Liu, J.D., Wen, W.E., Fu, L.P., Li, G.Y., Sui, X.X., Xia, X.C., Wang, C.P., Cao, S.H., 2017. Cloning of *TaTPP-6AL1* associated with grain weight in bread wheat and development of functional marker. Mol Breed. 37, 78.

Zheng, L., Karim, M.R., Hu, Y.G., Shen, R.F., Lan, P., 2021. Greater morphological and primary metabolic adaptations in roots contribute to phosphate-deficiency tolerance in the bread wheat cultivar Kenong199. BMC Plant Biol. 21, 381.

Zheng, J.L., Liu, G.H., Wang, S., Xia, G.M., Chen, T.T., Chen, Y.L., Siddique, K.H.M., Chi, D.C., 2022. Zeolite enhances phosphorus accumulation, translocation, and partitioning in rice under alternate wetting and drying. Field Crop Res. 286, 108632.

Zheng, L., Wang, R.N., Zhou, P.J., Pan, Y.L., Shen, R.F., Lan, P., 2023. Comparative physiological and proteomic response to phosphate deficiency between two wheat genotypes differing in phosphorus utilization efficiency. J Proteomics. 280, 104894.

Zhuang, M.J., Li, C.N., Wang, J.Y., Mao, X.G., Li, L., Yin, J., Du, Y., Wang, X., Jing, R.L., 2021. The wheat *SHORT ROOT LENGTH 1* gene *TaSRL1* controls root length in an auxin-dependent pathway. J Exp Bot. 72 (20), 6977–6989.

